## Supplemental Information for "A bead-based GPCR phosphorylation immunoassay for high-throughput ligand profiling and GRK inhibitor screening"

### Supplementary information

**This .pdf includes**

**Supplementary Figures: 15**

**Supplementary Tables: 6**

**Supplementary Figure 1**

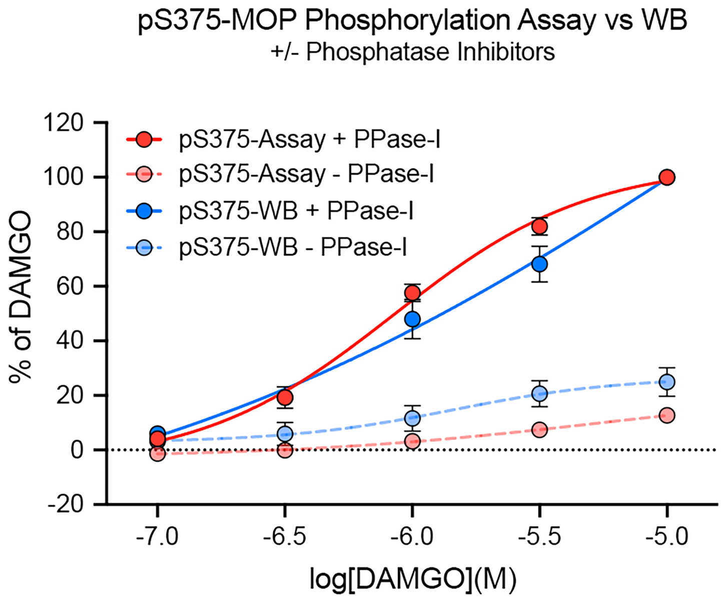


**Supplementary Figure 1.** Comparison of normalized pS375-MOP phosphorylation assay and western blot (WB) data in dephosphorylation experiments. MOP-HEK293 cells were treated with a DAMGO dilution series and lysed in detergent buffer in the presence or absence of protein phosphatase inhibitors (+/- PPase-I). Lysates were either processed according to 7TM phosphorylation assay (red) or western blot (blue) protocol. Graphs represent means of n=5 independent experiments performed in duplicates ± SEM. Western blot images were quantified using the ImageJ software. All data points were normalized to 10 µM DAMGO.

**Supplementary Figure 2**

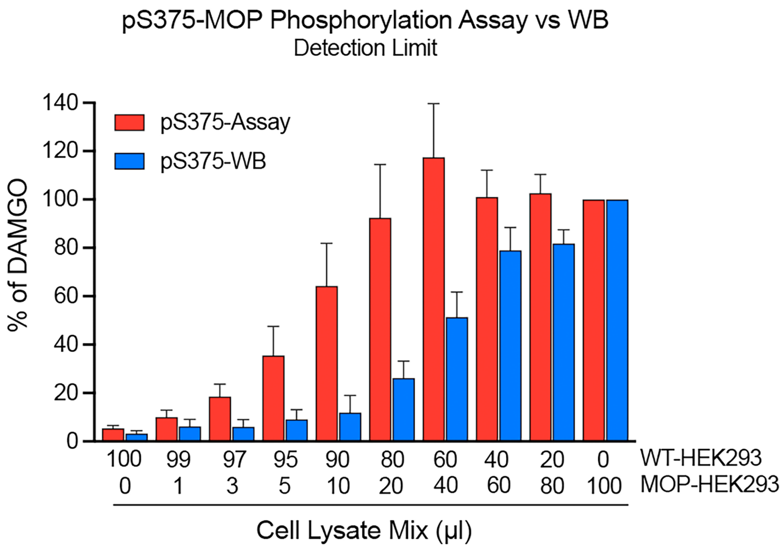


**Supplementary Figure 2.** Detection limits determined under 7TM phosphorylation assay and western blot (WB) conditions. Cell lysates from DAMGO-stimulated WT-HEK293 and MOP-HEK293 cells were mixed in different ratios to a final volume of 100 µl. They were then processed according to 7TM phosphorylation assay (red) or western blot (blue) protocol. Bars represent means of n=3 (western blot) to n=5 (assay) independent experiments performed in duplicates ± SEM. Western blot images were quantified using the ImageJ software. All data points are normalized to 100 µl DAMGO-stimulated MOP-HEK293 cell lysate.

**Supplementary Figure 3**

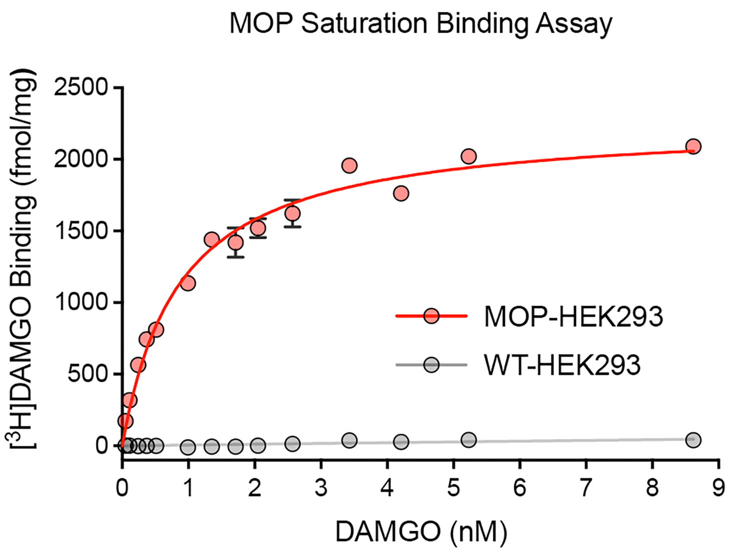


**Supplementary Figure 3.** Saturation binding of [$^3$H]DAMGO to membranes of MOP-HEK293 and WT-HEK293 cells. Increasing concentrations of [$^3$H]DAMGO were incubated with cell membranes at 25°C for 60 min in the absence or presence of 10 µM DAMGO as described in Methods. In MOP-HEK293 cells, the calculated $K_d$ value was 0.92 ± 0.07 nM and $B_{max}$ was 2279 ± 19 fmol/mg protein. Data are presented as means ± SEM of n = 3 independent experiments performed in duplicates.

**Supplementary Figure 4**

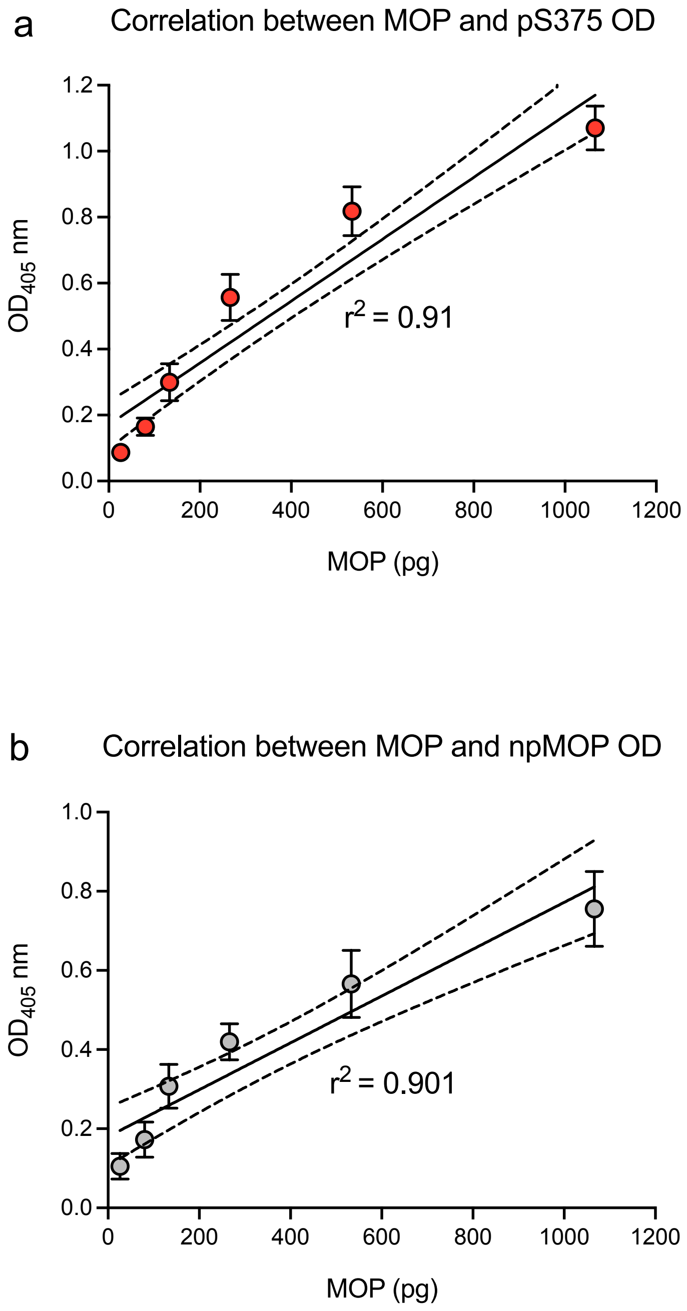


**Supplementary Figure 4.** Range of linear detection in 7TM phosphorylation assays. Cell lysates from DAMGO-stimulated WT-HEK293 and MOP-HEK293 cells were mixed in different ratios to a final volume of 100 µl. They were then processed for pS375-MOP (**a**) or np-MOP (**b**) according to 7TM phosphorylation assay. Data points represent means of n=5 independent experiments performed in duplicates ± SEM. Pearson correlation coefficients were calculated using GraphPad Prism.

**Supplementary Figure 5**

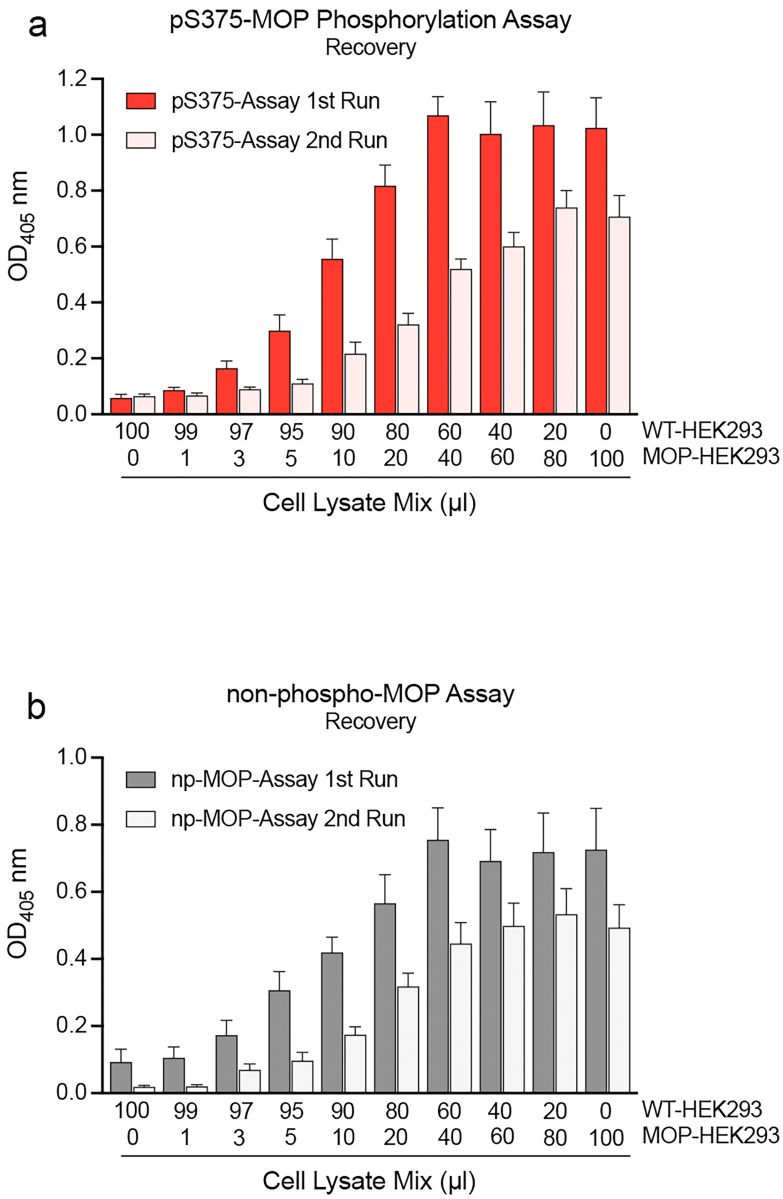


**Supplementary Figure 5.** OD signal intensity in recovery experiments. WT-HEK293 and MOP-HEK293 were treated with 10 µM DAMGO and combined in different ratios to yield 100 µl cell lysate mixtures. Lysates were incubated with anti-HA magnetic beads for receptor immunoprecipitation (1st run). Afterwards, the lysate-bead mix was placed on a magnet and the supernatant was transferred into a new 96-well plate with fresh magnetic beads. They were then incubated for another 2 h at 4°C (2nd run). Afterwards both plates were washed and further processed according to the 7TM phosphorylation assay protocol. OD signals were measured using pS375-MOP (**a**) or np-MOP (b). Datasets represent means of n=5 independent experiments performed in duplicates ± SEM.

**Supplementary Figure 6**

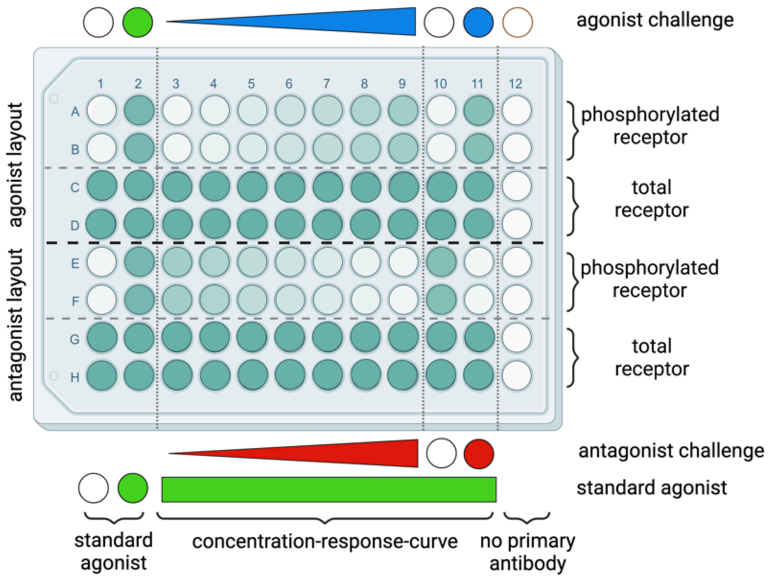


**Supplementary Figure 6.** Schematic model of the recommended 7TM phosphorylation assay setup to determine concentration-response-curves of agonists or antagonists. Two distinct compounds may be analysed in parallel on one 96-well plate allowing for determination of GPCR phosphorylation responses to different agonist or antagonist concentrations. Rows A to D may be used for the first, and rows E to H for the second compound. The compound dilution series is performed at least as duplicates in columns 1 to 9 ranging from 0 µM (unstimulated) to, for instance, 10 µM (fully saturated). If necessary, columns 10 and 11 are available for comparison to a known endogenous stimulant (agonist control). Every well used for detection of phosphorylation-specific signals (phosphorylated receptor) has a corresponding well as a loading control (total receptor) (e.g. A1 and C1). No primary antibody is added to column 12 (negative control, background signal). (Created with BioRender.com)

**Supplementary Figure 7**

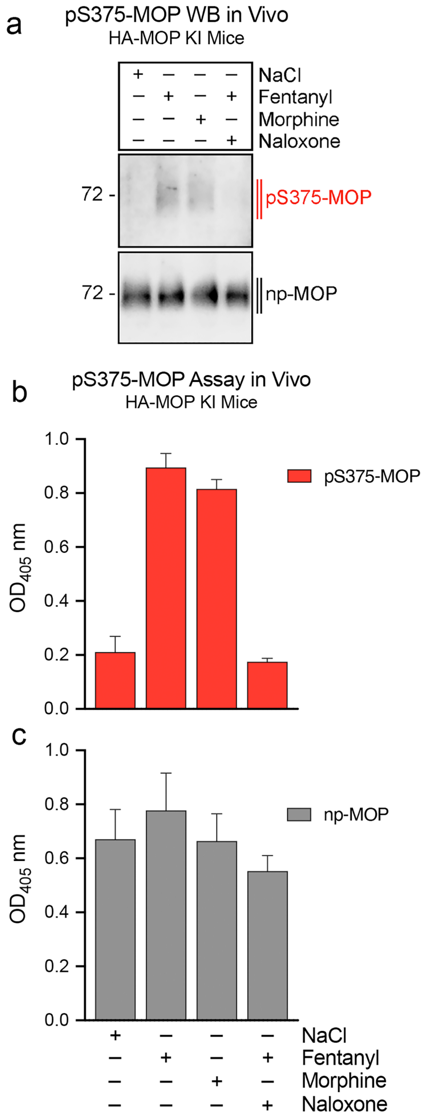


**Supplementary Figure 7.** Application of the 7TM phosphorylation assay for analysis of *in vivo* samples. HA-MOP KI mice were injected subcutaneously with either NaCl (0.9%, 30 min), fentanyl (0.3 mg/kg, 15 min), morphine (30 mg/kg, 30 min) or naloxone (2 mg/kg, 10 min) and fentanyl (0.3 mg/kg, 15 min). Mouse brains were then isolated and homogenized before adding anti-HA magnetic beads. Samples were processed according to western blot (**a**) or 7TM phosphorylation assay (**b**, **c**) protocol. Western blot images are representatives of n=3 independent experiments. OD signals are displayed as mean of n=5 independent assay replicates ± SEM.

### Supplementary Figure 8

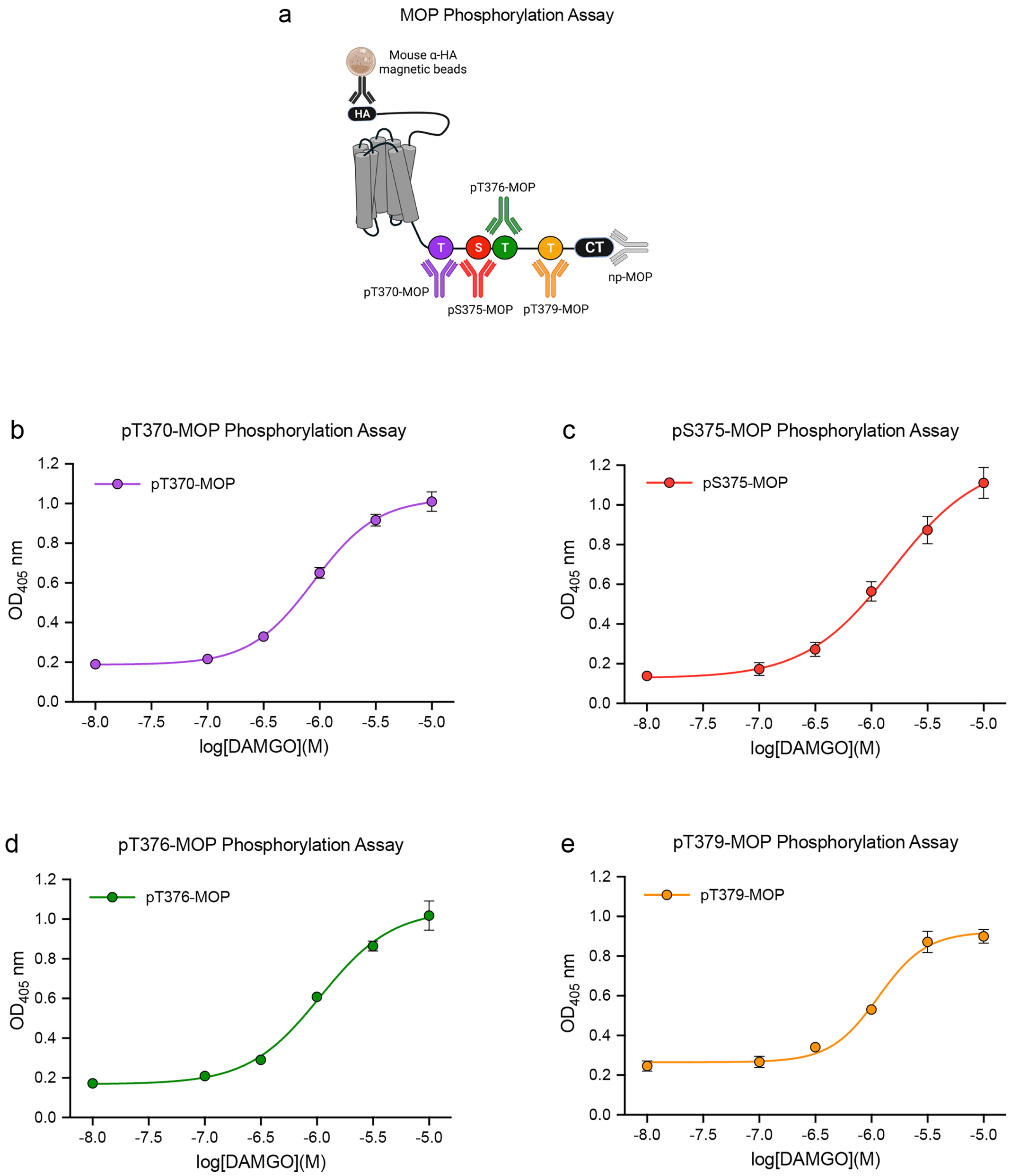


**Supplementary Figure 8.** OD signals in MOP phosphorylation assays. (**a**) Schematic representation of MOP depicting antibody binding sites for pT370-, pS375-, pT376-, pT379- and np-MOP. (**b-c**) MOP-HEK293 cells were treated with increasing DAMGO concentrations. Phosphorylation was determined using pT370-MOP (**b**), pS375-MOP (**c**), pT376-MOP (**d**) and pT379-MOP (**e**) antibodies according to 7TM phosphorylation assay protocol. All data points represent means of n=5 independent experiments performed in duplicates ± SEM.

**Supplementary Figure 9**

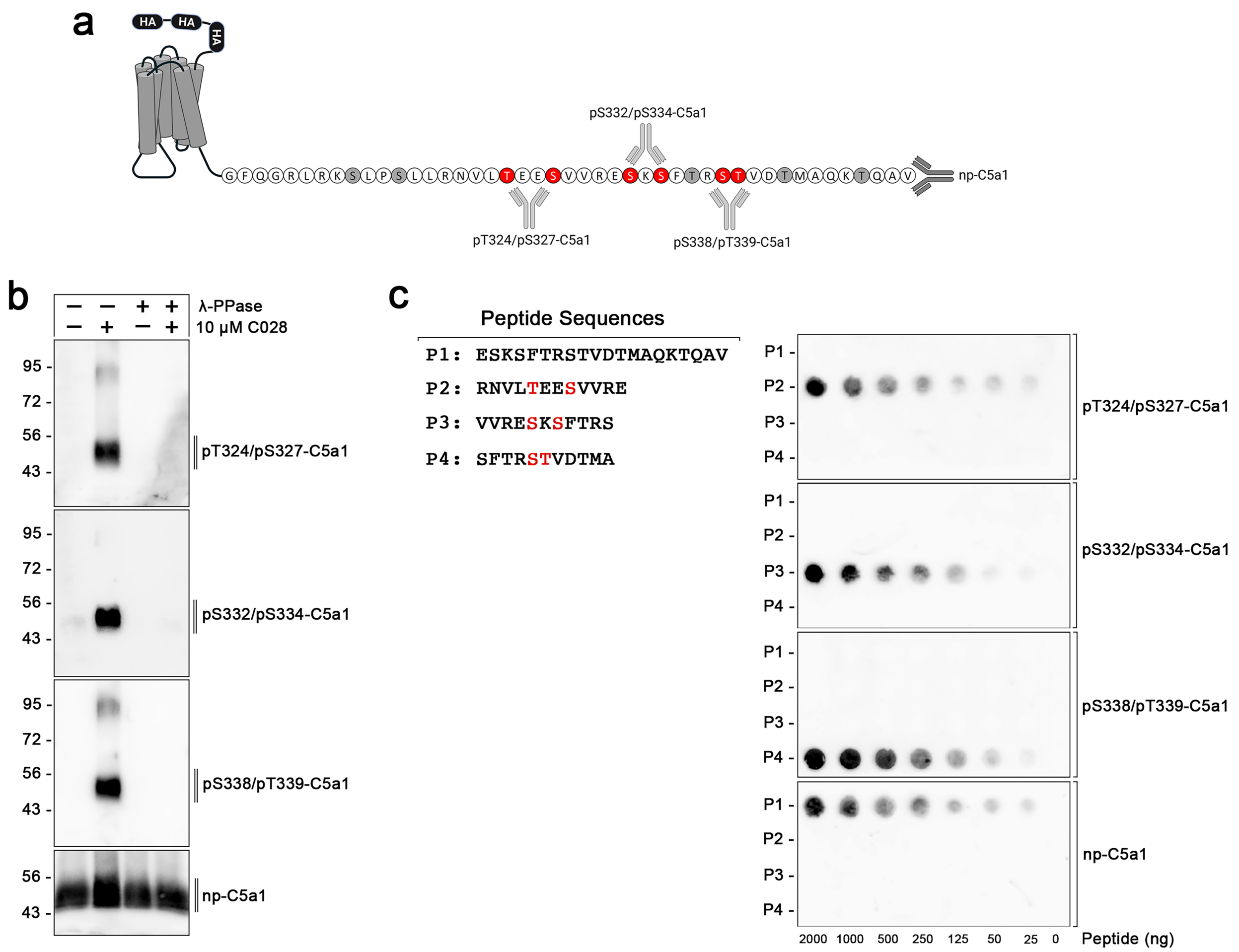


**Supplementary Figure 9.** Characterization of phosphor-C5a1 antibodies. (**a**) Schematic representation of C5a1 depicting antibody binding sites for pT324/pS327-, pS332/pS334-, pS338/pT339- and np-C5a1. S/T sites detected using phosphosites-specific C5a1 antibodies are depicted in red. All other potential phosphate acceptor sites are depicted in gray. (**b**) C5aR1-HEK293 cells were either left unstimulated or stimulated with 10 µM C028 (+). During immunoprecipitation of the receptor, samples were additionally treated in presence (+) or absence (-) of λ-protein phosphatase (λ-PPase). Western blot analysis shows phosphorylation at T324/S327-, S332/S334- and S338/T339-C5aR1. The np-C5aR1 antibody ensures equal receptor loading in every sample. (**c**) A dilution series of the peptides P1-P4 were applied onto a PVDF membrane. Dot blot analysis depicts pT324/pS327-, pS332/pS334-, pS338/pT339- and np-C5a1 antibody binding. Note that pT324/pS327-, pS332/pS334-, pS338/pT339- and np-C5a1 antibodies detect their cognate target without detectable cross-reactivity. (**b**) and (**c**) display a representative of n=4 independent experiments.

### Supplementary Figure 10

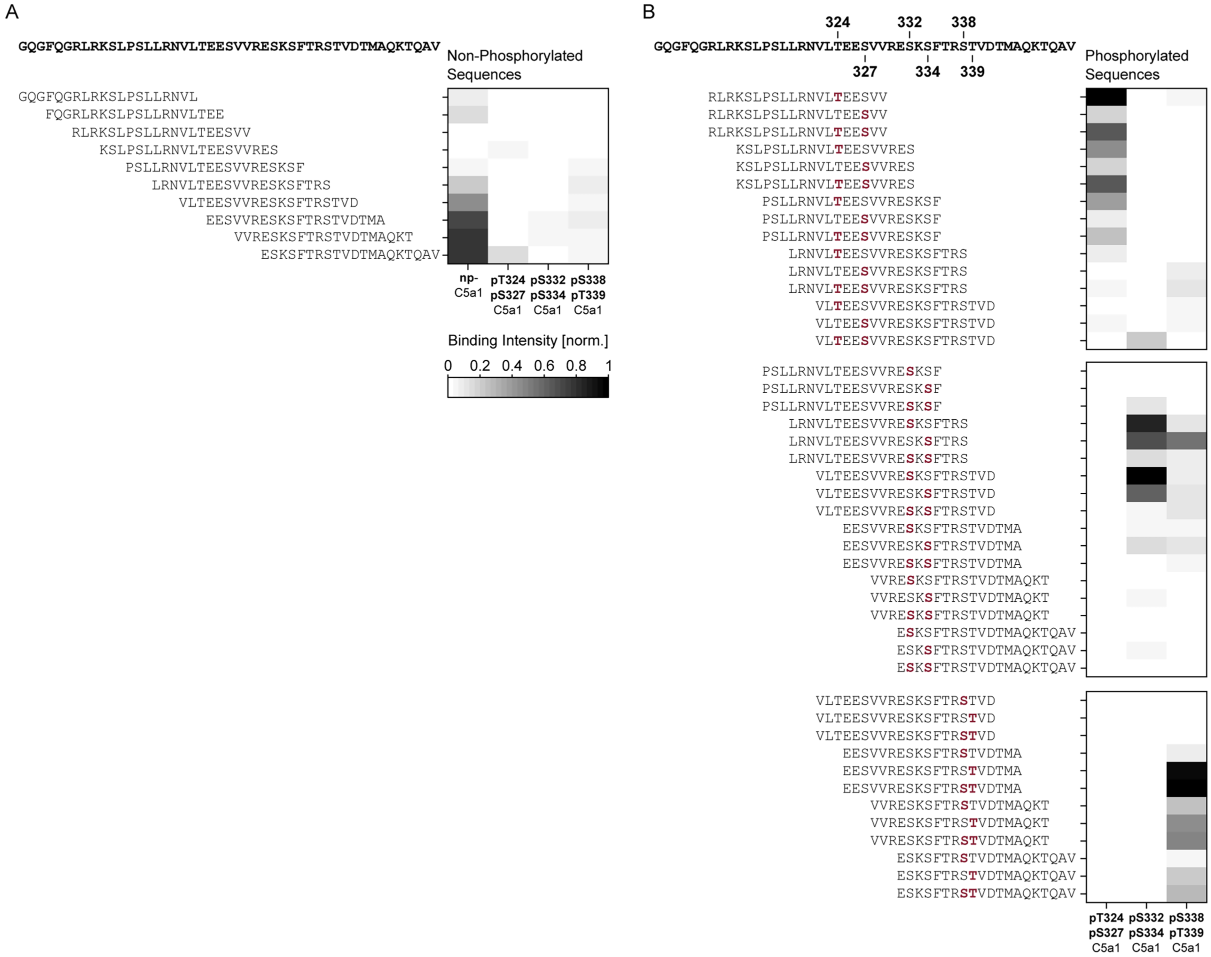


**Supplementary Figure 10.** Microarray based mapping of pT324/pS327-, pS332/pS334-, pS338/pT339- and np-C5a1 antibodies. An overlapping peptide library (peptide length 15 and offset of 3 amino acids), containing phosphorylated serine and threonine building blocks was used to validate the phosphosite specificity. Note that pT324/pS327-, pS332/pS334-, pS338/pT339- and np-C5a1 antibodies detect their cognate target with minimal detectable cross-reactivity. All data points represent means of n=3 independent experiments.

**Supplementary Figure 11**

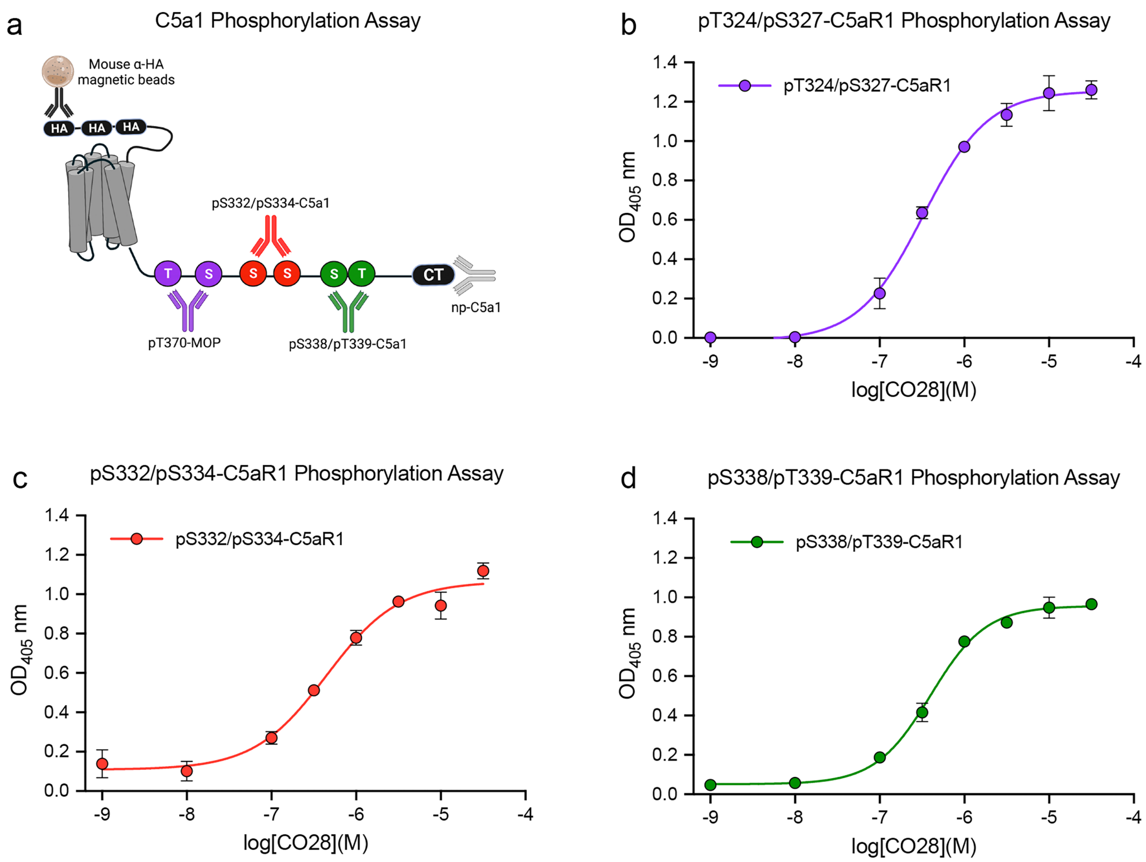


**Supplementary Figure 11.** OD signals in C5a1 phosphorylation assays. (**a**) Schematic representation of C5a1 depicting antibody binding sites for pT324/pS327-, pS332/pS334-, pS338/pT339- and np-C5a1. (**b-c**) C5a1-HEK293 cells were treated with increasing C028 concentrations. Phosphorylation was determined using pT324/pS327-C5a1 (**b**), pS332/pS334-C5a1 (**c**) and pS338/pT339-C5a1 (**d**) antibodies according to 7TM phosphorylation assay protocol. All data points represent means of n=5 independent experiments performed in duplicates ± SEM.

**Supplementary Figure 12**

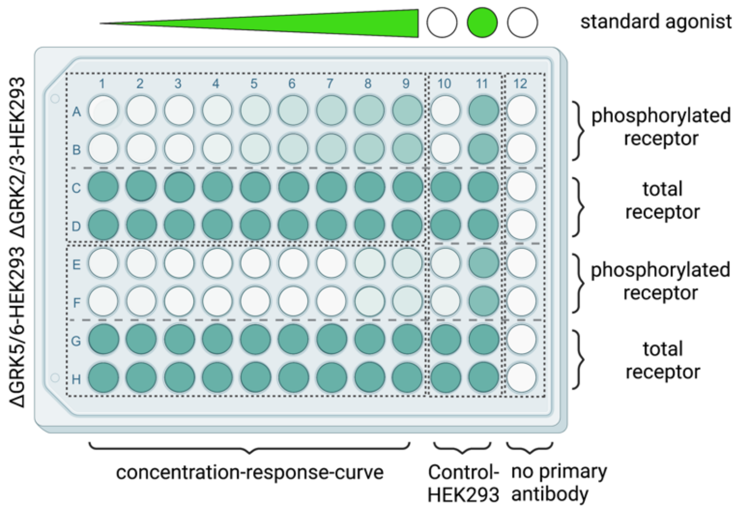


**Supplementary Figure 12.** Recommended layout of the 7TM phosphorylation assay for experiments with ΔGRK-HEK293 cells. In the 96-well format, rows A to D may be used for the first, and rows E to H for the second GRK KO cell line. Columns 1 to 9 encompass a dilution series of the agonist of interest (green). Columns 10 and 11 are used for the comparison with unstimulated and stimulated control-HEK293 cells. Every well used for the detection of phosphorylation-specific signals (phosphorylated receptor) has a corresponding well for the loading control by using phosphorylation-unspecific antibodies (total receptor) (e.g. A1 corresponds to C1). No primary antibody is added to column 12 (background signal). The consecutive treatment of all wells remains constant. Background signal, loading control as well as unstimulated and stimulated control-HEK293 signals are included in the calculation of concentration-response curves. (Created with BioRender.com)

**Supplementary Figure 13**

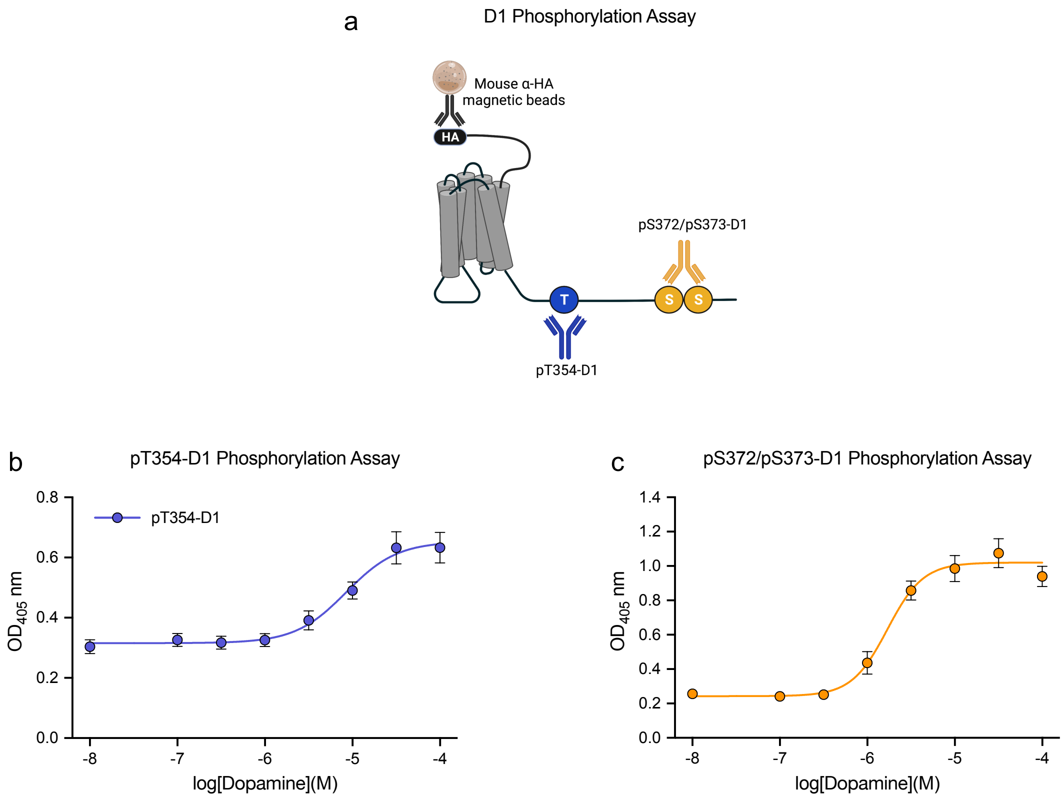


**Supplementary Figure 13.** OD signals in D1 dopamine receptor phosphorylation assays. (**a**) Schematic representation of D1 dopamine receptor depicting antibody binding sites for pT354- and pS372/pS373-D1. (**b**, **c**) D1-HEK293 cells were treated with increasing dopamine concentrations. Phosphorylation was determined using pT354-D1 (**b**) and pS372/pS373-D1 (**c**) antibodies according to 7TM phosphorylation assay protocol. All data points represent means of n=4 independent experiments performed in duplicates ± SEM.

### Supplementary Figure 14

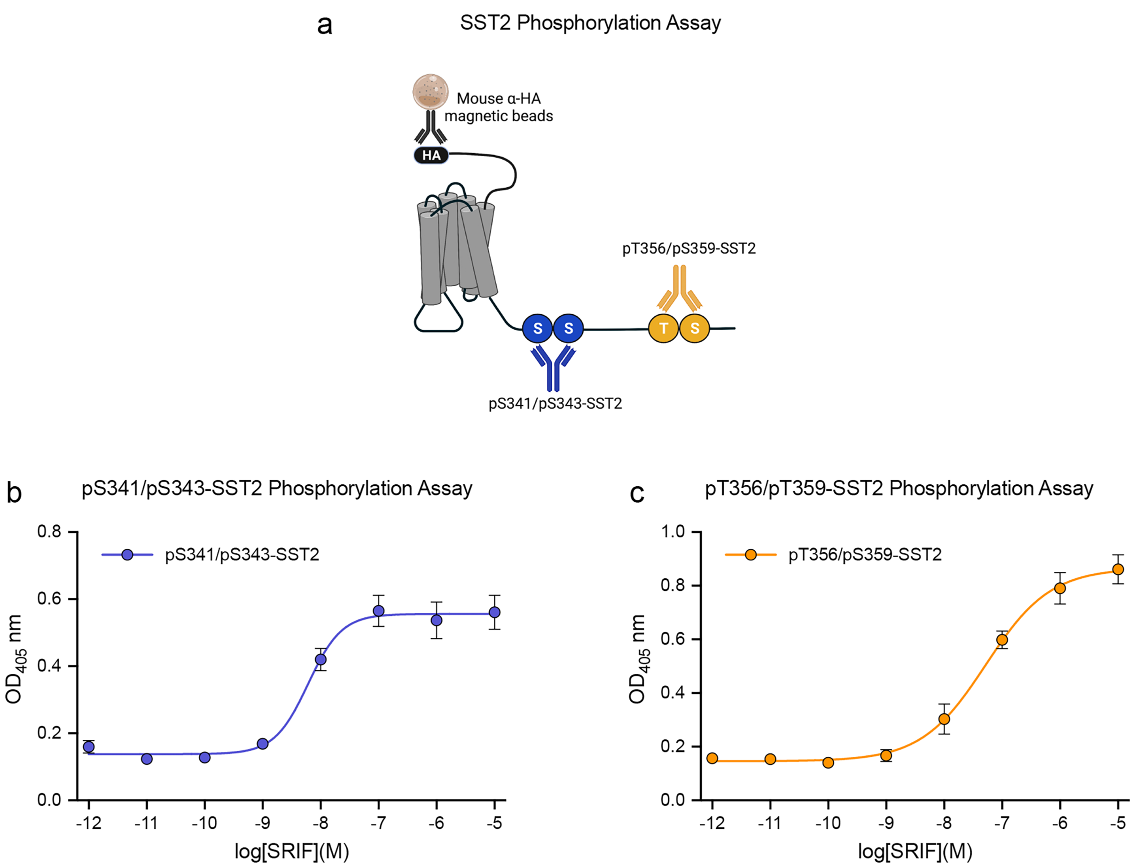


**Supplementary Figure 14.** OD signals in SST2 somatostatin receptor phosphorylation assays. (**a**) Schematic representation of SST2 receptor depicting antibody binding sites for pS341/pS343- and pT356/pT359-SST2. (**b**, **c**) SST2-HEK293 cells were treated with increasing SRIF concentrations. Phosphorylation was determined using pS341/pS343-SST2 (**b**) and pT356/pT359-SST2 (**c**) antibodies according to 7TM phosphorylation assay protocol. All data points represent means of n=4 independent experiments performed in duplicates ± SEM.

**Supplementary Figure 15**

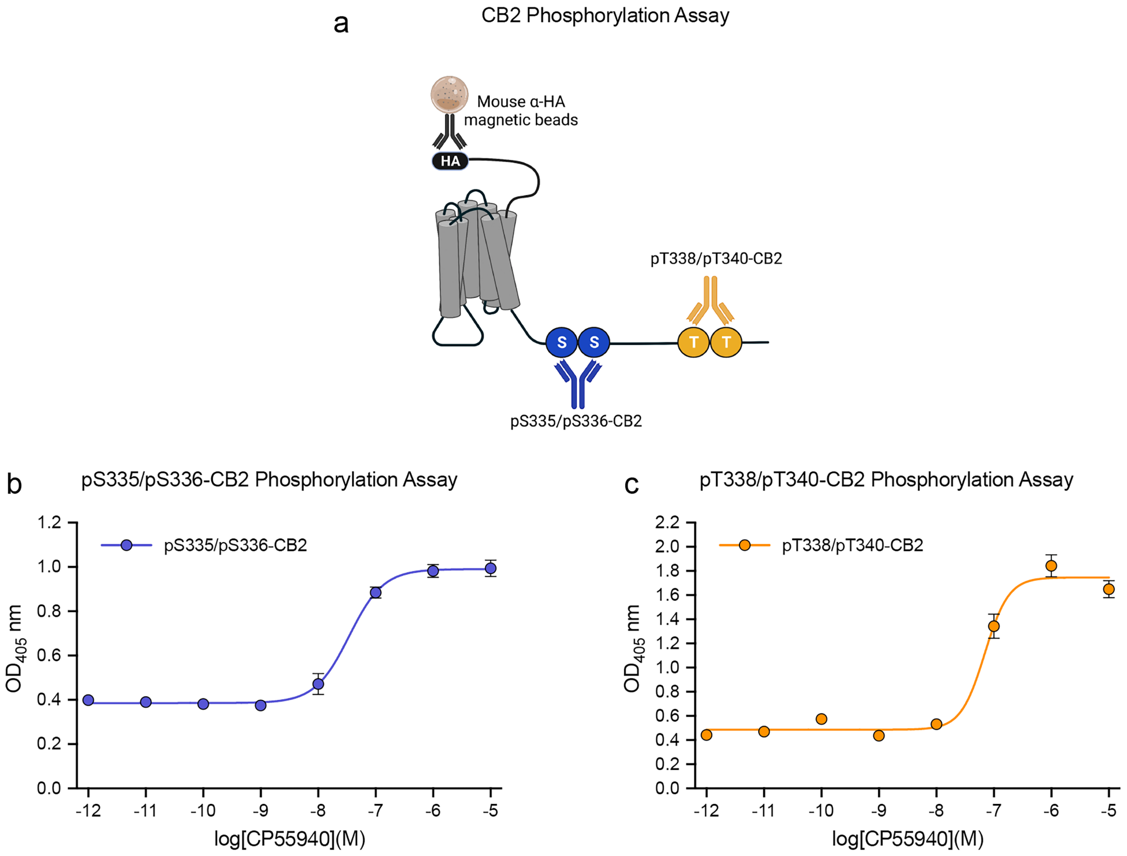


**Supplementary Figure 15.** OD signals in CB2 cannabinoid receptor phosphorylation assays. (**a**) Schematic representation of CB2 receptor depicting antibody binding sites for pS335/pS336- and pT338/pT339-CB2. (**b**, **c**) CB2-HEK293 cells were treated with increasing CP55940 concentrations. Phosphorylation was determined using pS335/pS336-CB2 (**b**) and pT338/pT339-CB2 (**c**) antibodies according to 7TM phosphorylation assay protocol. All data points represent means of n=4 independent experiments performed in duplicates ± SEM.

**Supplementary Table 1**. Comparison of agonist-induced MOP phosphorylation, G-protein signaling, and GRK and arrestin binding

| | DAMGO | Morphine |
|---|---|---|
| Assay | $pEC_{50}$ | $pEC_{50}$ |
| p-MOP | 6.0 ± 0.02 | 5.4 ± 0.21 |
| GRK2 | 6.6 ± 0.19 | 6.8 ± 0.17 |
| GRK3 | 6.4 ± 0.17 | - |
| β-Arrestin1 | 5.9 ± 0.12 | - |
| β-Arrestin2 | 6.0 ± 0.02 | - |
| GIRK | 8.9 ± 0.03 | 8.30 ± 0.05 |

Data represent the mean of at least n=3 independent experiments performed in duplicates ± SE.

**Supplementary Table 2**. Differential involvement of GRK2/3 and GRK5/6 in C5a1 multisite phosphorylation

| | Control-HEK293 | | ΔGRK2/3-HEK293 | | ΔGRK5/6-HEK293 | |
|---|---|---|---|---|---|---|
| Phosphorylation Assay | $pEC_{50}$ | $E_{max}$ (% Control C028) | $pEC_{50}$ | $E_{max}$ (% Control C028) | $pEC_{50}$ | $E_{max}$ (% Control C028) |
| pT324/pS327-C5a1 | 6.3 ± 0.13 | 100 ± 3.5 | 6.2 ± 0.09 | 79.8 ± 2.5 | - | 27.8 ± 3.7 |
| pS332/pS334-C5a1 | 6.4 ± 0.21 | 100 ± 5.3 | 6.3 ± 0.13 | 70.2 ± 2.1 | - | 50.82 ± 8.7 |
| pS338/pT339-C5a1 | 6.2 ± 0.07 | 100 ± 2.4 | 6.0 ± 0.19 | 55.2 ± 9.5 | - | 63.90 ± 8.8 |

Data represent the mean of n=5 independent experiments performed in duplicates ± SE.

**Supplementary Table 3**. In vitro GRK inhibitor profiling by Lance kinase activity assay

| | GRK2 Assay | GRK5 Assay |
|---|---|---|
| Inhibitor | $IC_{50}$ (nM) | $IC_{50}$ (nM) |
| LDC9728 | 39 ± 13 | 10 ± 4 |
| LDC8988 | 1519 ± 213 | 14 ± 2 |
| Compound101 | 35 ± 12 | >10,000 |

Data represent the mean of n=3 independent experiments ± SE.

**Supplementary Table 4.** 7TM phosphorylation assay parameters

| Receptor | Assay | Agonist | $pEC_{50}$ | $OD_{405}$ min | $OD_{405}$ max |
|---|---|---|---|---|---|
| **MOP** | pT370-MOP | DAMGO | 6.1 ± 0.04 | 0.19 | 1.03 |
| | pS375-MOP | DAMGO | 6.0 ± 0.05 | 0.13 | 1.23 |
| | pT376-MOP | DAMGO | 6.0 ± 0.03 | 0.17 | 1.06 |
| | pT379-MOP | DAMGO | 6.0 ± 0.06 | 0.26 | 0.93 |
| **C5a1** | pT324/pS327-C5a1 | C028 | 6.3 ± 0.13 | 0.01 | 1.26 |
| | pS332/pS334-C5a1 | C028 | 6.4 ± 0.21 | 0.11 | 1.06 |
| | pS338/pT339-C5a1 | C028 | 6.2 ± 0.07 | 0.05 | 0.97 |
| **D1** | pT354-D1 | Dopamine | 5.1 ± 0.14 | 0.31 | 0.65 |
| | pS372/pS373-D1 | Dopamine | 5.8 ± 0.07 | 0.24 | 1.02 |
| **SST2** | pS341/pS342-SST2 | SRIF | 8.2 ± 0.14 | 0.14 | 0.56 |
| | pT356/pT359-SST2 | SRIF | 7.3 ± 0.13 | 0.15 | 0.86 |
| **CB2** | pS335/pS336-CB2 | CP55940 | 7.5 ± 0.08 | 0.38 | 0.99 |
| | pT338/pT339-CB2 | CP55940 | 7.2 ± 0.11 | 0.48 | 1.74 |

Data represent the mean of n=4-6 independent experiments performed in duplicates ± SE.

**Supplementary Table 5:** Reagents and concentrations in GRK2 kinase assay

| Reagents | Stock solution | Working solution | Final assay | Supplier |
|---|---|---|---|---|
| Ulight-peptide substrate | 5 µM | 125 nM | 50 nM | PerkinElmer |
| peptide antibody | 625 nM | 4nM | 2 nM | PerkinElmer |
| GRK2 | 5.95 µM | 50 nM | 20 nM | ThermoFisher |
| ATP low | 100 mM | 75 µM | 15 µM | Sigma |

**Supplementary Table 6:** Reagents and concentrations in GRK5 kinase assay

| Reagents | Stock solution | Working solution | Final assay | Supplier |
|---|---|---|---|---|
| Ulight-peptide substrate | 5 µM | 125 nM | 50 nM | PerkinElmer |
| peptide antibody | 625 nM | 4nM | 2 nM | PerkinElmer |
| GRK5 | 12.73 µM | 25 nM | 10 nM | ThermoFisher |
| ATP low | 100 mM | 32.5 µM | 6.5 µM | Sigma |
